# Amplicon sequencing reveals complex infection in infants congenitally infected with *Trypanosoma cruzi* and informs the dynamics of parasite transmission

**DOI:** 10.1101/2022.11.16.516746

**Authors:** Jill Hakim, Andreea Waltmann, Freddy Tinajeros, Oksana Kharabora, Edith Málaga Machaca, Maritza Calderon, María del Carmen Menduiña, Jeremy Wang, Daniel Rueda, Mirko Zimic, Manuela Verástegui, Jonathan J Juliano, Robert H Gilman, Monica R. Mugnier, Natalie M Bowman, Chagas working group

**Author notes:** co-senior author (Monica R. Mugnier, Natalie M Bowman).

## Abstract

Congenital transmission of *Trypanosoma cruzi*, the causative agent of Chagas disease, is an important source of new infections worldwide. The mechanisms of congenital transmission remain poorly understood, but there is evidence that parasite factors could play a role.

Investigating changes in parasite strain diversity during transmission could provide insight into the parasite factors that influence the process. Here we use deep amplicon sequencing of a single copy gene in the *T. cruzi* genome to evaluate the diversity of infection in a collection of clinical blood samples from Chagas positive mothers and their infected infants. We found several infants and mothers infected with more than two parasite haplotypes, indicating infection with multiple parasite strains. Two haplotypes were detected exclusively in infant samples, while one haplotype was never found in infants, suggesting a relationship between the probability of transmission and parasite genotype. Finally, we found an increase in parasite population diversity in children after birth compared to their mothers, suggesting that there is no transmission bottleneck during congenital infection and that multiple parasites breach the placenta in the course of congenital transmission.

## Background

*Trypanosoma cruzi* is the causative agent of Chagas disease and is estimated to infect almost 6 million people worldwide (1). Effective vector control has decreased the number of new infections, but congenital transmission has become an increasing concern, particularly in non-endemic areas. An estimated 22% of new Chagas cases occur via congenital transmission, and approximately 5% of *T. cruzi*-infected mothers will transmit the parasite to their newborn (1,2). The role that parasite genetics plays in transmission and congenital infection is poorly understood. Identifying parasite strains that are more likely to be vertically transmitted could uncover mechanisms underlying this growing source of new cases and lead to improved detection and treatment of congenital *T. cruzi* infection.

Studies examining parasite diversity within *T. cruzi* infections are often performed with a focus on *T. cruzi’s* six Discrete Typing Units (DTUs), TcI through TcVI, which are distinct genetic groups that segregate by genotyped markers. However, diversity has been observed within individual DTUs, both across isolates and within single infections (3–5). Therefore, using DTUs alone likely underestimates parasite diversity, as several clones of the same DTU could co-infect a patient (6,7). Moreover, *T. cruzi* has two hybrid DTUs, TcV and TcVI. Each of these are ancient hybrids of TcII and TcIII, with each homologous chromosome of the parasite thought to approximately match one of these ancestral parental haplotypes. In some parasite clones, there may be complex recombination events present between these parental haplotypes resulting in mosaic alleles (8). Diversity arising from these events can only be identified by characterization of each individual haplotype. Whole genome sequencing of clinical isolates can circumvent this problem and identify complex hybrids but is typically not feasible due to the low parasitemia during chronic infection. To ameliorate the problem of low parasitemia, some studies have targeted high copy number genes, such as the miniexon locus, to evaluate the complexity of *T. cruzi* infection (9–11). These genes frequently display variability even within a single parasite strain, however, artificially raising the apparent number of parasite clones (3,12).

Here, we use amplicon sequencing of a single copy gene, *TcSC5D*, to characterize the clonal diversity in clinical samples of Chagas positive infants and mothers, including several twins. Importantly, this gene contains nucleotide polymorphisms that are distinct across several DTUs, allowing an additional rough DTU determination (13). Our results reveal haplotypes that are present exclusively in infant or mother samples, indicating a possible relationship between parasite genotype and transmission. We also observed an increase in parasite diversity in infant samples relative to maternal samples, suggesting that there is no bottleneck during congenital transmission of *T. cruzi* and that transmission may be the result of multiple colonizing parasites that infect the infant during pregnancy.

## Methods

### Study information

Chagas positive mothers were recruited from Percy Boland Women’s Hospital in Santa Cruz, Bolivia between the years of 2016 and 2018. 300 µl of maternal venous blood and 300 µl of infant blood from heel puncture was taken at birth. For longitudinal timepoints at one, three, and nine months, 300 µl of infant venous blood was taken. Mothers recruited to the study were surveyed regarding obstetrics and demographic characteristics. Analysis of these epidemiological characteristics are described elsewhere (14), and epidemiological data from the patients involved in this study are provided in supplemental data (supplemental table 1). This collection protocol was approved by the ethics committee of the Bolivian Catholic University, international registration FWA 0017928 and PRISMA 00001219. Study analysis was exempted by the Institutional Review Board at the University of North Carolina at Chapel Hill (IRB 19–3014).

### Sample Processing and Amplicon Sequencing

DNA was extracted at the Infectious Diseases Research Laboratory of the Universidad Peruana Cayetano Heredia in Lima, Peru. Amplicon PCR, library prep and sequencing were performed at University of North Carolina at Chapel Hill. Details of nested PCR and library preparation are described in supplementary methods. Raw sequencing data are available in National Center for Biotechnology Information (NCBI) Sequence Read Archive under accession number PRJNA891347.

### Haplotype calling

Demultiplexed reads were adapter-trimmed using CutAdapt (15). Following trimming, any read pairs less than 100 base pairs long were removed to eliminate adapter dimers and off-target amplicons from the pipeline. Following adapter trimming, reads were aligned to a custom BLAST search against the target amplicon to remove non-specific reads and orient amplicons in the same direction prior to calling haplotypes. Filtered reads were then run through DADA2 using a max expected error of 2 for both the R1 and R2 reads (16). Any amplicon sequence variants that made up less than 0.01% of the total population of reads were removed from the analysis, and any sample that had fewer than 200 total reads after all filtering steps was removed from the analysis.

Two of the recovered haplotypes were identical, save for a small insertion at the end of the amplicon. Because it is possible that the shorter amplicon could be amplified in samples containing the longer amplicon, they were merged into one amplicon for the analysis. This amplicon sequence is available in the supplemental data as haplotype number 12, and it was merged with haplotype 0 for subsequent analysis.

### Haplotype diversity analysis

Parasite diversity was measured using Shannon’s diversity index using the Vegan R package (17). Only haplotypes detected in both sample replicates were counted when analyzing the frequency of appearance of each haplotype, except in cases where only one sample replicate passed filtering; in these cases, every detected haplotype was counted. All scripts used to process raw data, call haplotypes, and generate figures for analysis are available at https://github.com/MugnierLab/Hakim2022.

## Results

### Haplotype sequences were largely similar and mostly belonged to hybrid DTUs TcV or TcVI

PCR targeting the *TcSC5D* gene was performed in duplicate on 75 total samples. After eliminating samples with low quality sequencing data, 44 samples were included in the analysis. These 44 samples included eight mother-infant sets (2 sets of twins, 6 singletons), 2 mothers without a matching infant sample, and 10 infants (including one set of twins) without a matching mother sample. All collected epidemiological data associated with these samples is available in supplemental table 1, alongside the total number of reads each sample replicate received.

The twelve unique haplotype sequences identified in the study population showed between 90.9% and 99.8% sequence similarity (Fig 1a). Using reference whole genome data as well as Sanger sequencing data at the same locus as a reference, we assigned each recovered haplotype a tentative DTU designation. DTU TcVI is an ancestral hybrid of a TcIII and TcII strain, and both Esmeraldo (resembling the TcIII ancestor) and non-Esmeraldo (resembling the TcII ancestor) homologous chromosomes were included as reference haplotypes. Figure 1b shows a table of all single nucleotide polymorphisms (SNPs) across the recovered haplotypes and reference sequences. Haplotype 6 shared 100% sequence identity to the TcI Sylvio strain, while all other haplotypes have some similarity to phased haplotypes of the hybrid CL Brener strain. Haplotypes 3, 8, and 11 had unique SNPs not found in the reference strains or in any other haplotype. Several haplotypes had stretches of similarity matching the TcII SNP pattern, before switching to match the TcIII pattern, and vice versa. Similarly, haplotypes 0, 3, and 8 have loci that suggest they may belong to TcII, but at site 82 all three have a nucleotide associated with TcIII. These patterns are consistent with recombination between homologous chromosomes in the hybrid strains. Though we cannot call these haplotypes’ DTU designation specifically, it is apparent that they are all hybrids, making them either TcV or TcVI.

**Figure 1:**
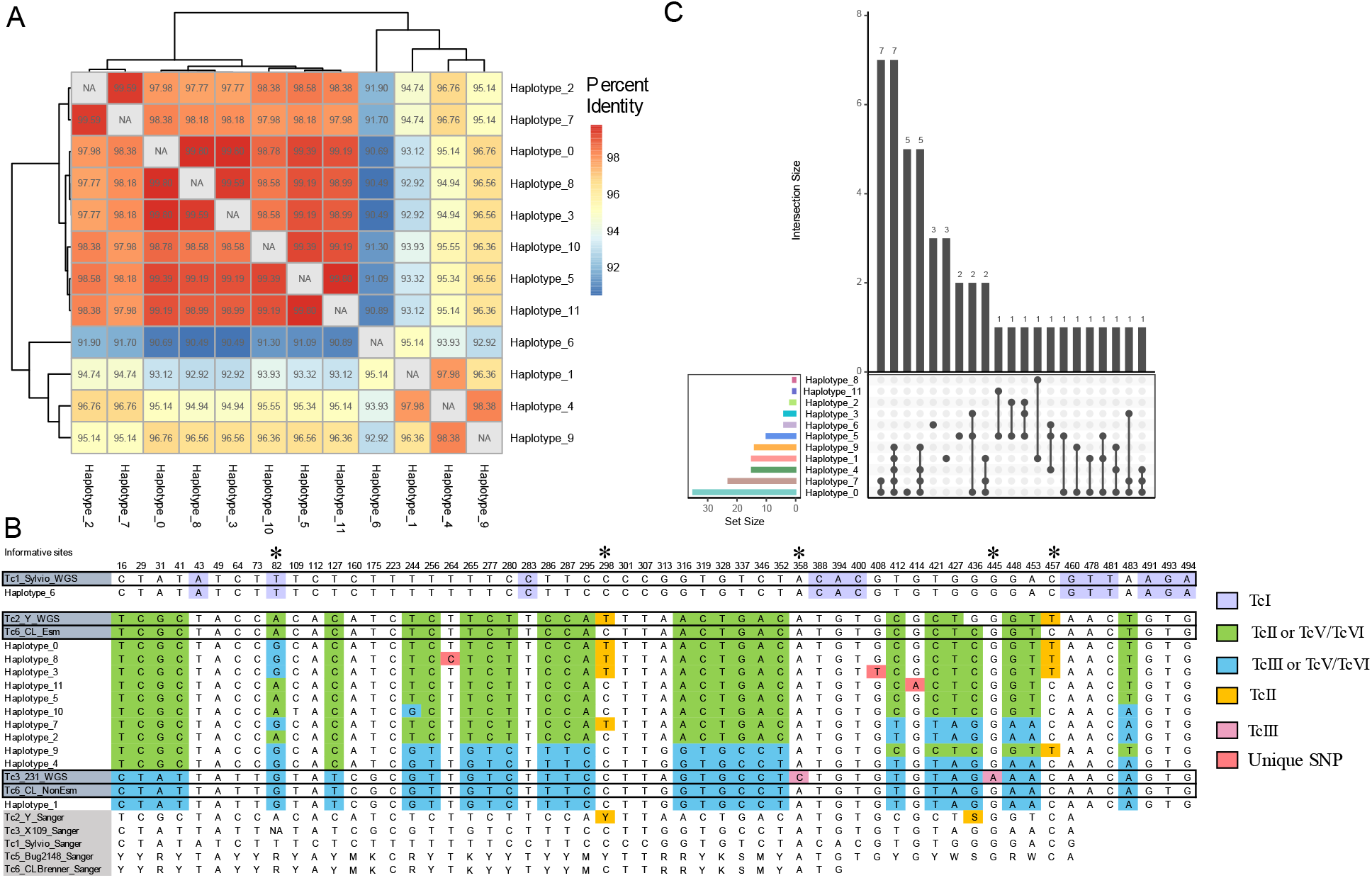
Many haplotypes belonged to the same DTU. **A**. Heatmap showing percent identity between haplotypes, clustered by similarity. **B** Table of all SNPs across haplotypes found. The position of each polymorphism within the amplicon is noted at the top of each column. Whole genome data of TcI, TcII, TcIII, and TcVI as well as Sanger sequencing data for a portion of the amplicon from TcI, TcII, TcIII, TcV and TcVI are included for reference. Bases are colored based on similarity to reference DTU data; only positions that are unique to a reference haplotype are colored. Asterixis denote informative SNP sites for DTU calls. **C** UpSet plot describing the frequency and co-occurrence of each haplotype and haplotype combination. Dots represent a haplotype or haplotype combination, with the frequency of that combination represented on the bar graph above.

We analyzed the combinations of haplotypes detected in each infection in order to determine whether certain haplotypes were more likely to co-occur or occur alone. Four haplotypes, haplotypes 0, 1, 5, and 6, were detected with no other haplotypes in some infections (Fig 1b). This could indicate that these haplotypes represent individual parasite clones in which both homologous chromosomes contain the same sequence at the amplicon locus. Most haplotypes occur in multiple infections, suggesting that the haplotypes found in infants represent parasite clones that were transmitted to the infant from the mother, rather than novel mutations arising during infection or transmission (Fig 1c). The analysis revealed 11 samples containing three or more haplotypes, which, because of *T. cruzi’s* diploid genome, likely indicates infection with at least two parasite clones. All of these samples were infant samples, with no complex infections detected in any of the 14 maternal samples (Supplemental figure 1). This finding should be interpreted with caution, however, as we used a stringent approach for haplotype counting in which haplotypes not occurring in the replicate sample are eliminated.

### Parasite strain diversity is higher in infants than in mothers

Given the presence of multiple parasite clones in many samples, we assessed the changes in parasite diversity that occur during congenital transmission. Under many modes of infectious transmission, a bottleneck occurs, and the colonized site has less strain diversity than the source. However, we find that maternal samples are less diverse, as measured by Shannon’s diversity, than samples of infants at birth (Fig 2a). Shannon’s diversity is more robust to sampling error than comparing the number of detected haplotypes, because it considers the proportion at which each haplotype is found(18). Diversity appears to gradually decrease as infants get older, though this effect is not statistically significant. A potential explanation for this observation could be that the low parasitemia in the maternal blood sample causes under sampling of the true maternal diversity of circulating parasites. We found that the parasitemia of each patient as measured by PCR was correlated to the Shannon’s diversity for each sample (Fig 2b). The effect was significant in the overall comparison (Spearman’s rho = -0.62, p =2.3e-4), but when stratifying by sample type, the maternal samples were not significantly correlated (Spearman’s rho = -0.14, p =0.75), suggesting that the maternal samples were sufficiently sampled while diversity in the infant samples may be underestimated. However, because there were no chronically infected maternal samples of a high enough parasitemia to directly compare to the acutely infected infant samples, and because there may be a causative link between blood parasite load and parasite diversity, it is impossible to eliminate the role that sampling may play in the reduced parasite diversity found in mother’s blood.

**Figure 2:**
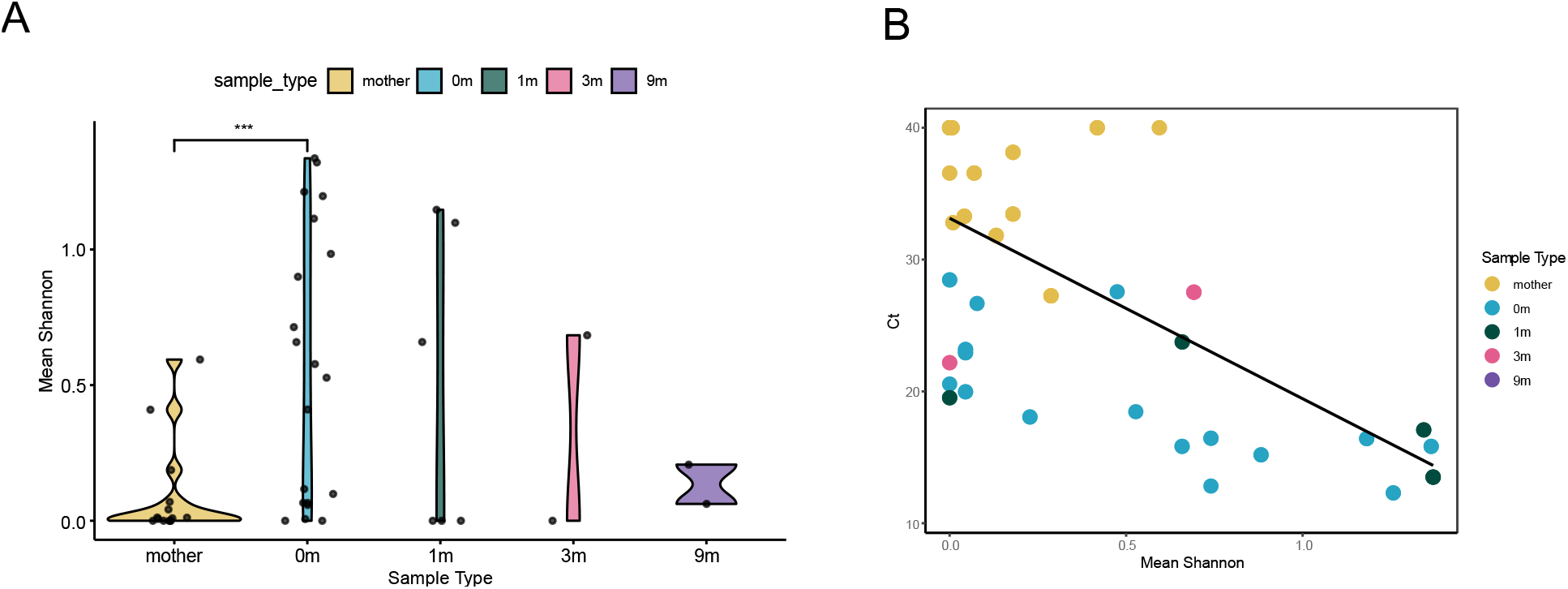
Haplotype diversity in mothers is lower than in infants at birth. **A**. Mean Shannon’s diversity between sample replicates for each sample type. 0m is infant’s sample at birth, 1m = one month of age, 3m = three months, 9m = nine months. Wilcoxon’s Rank sum test, p = 0.0024. **B**. Association between parasitemia measured by CT and average Shannon’s diversity for each replicate, with undetermined samples set at CT = 40. Spearman’s correlation was done excluding samples with undetermined CT. Mother’s samples: Spearman’s rho = -0.14, p = 0.752. 0m samples: Spearman’s rho = -0.82, p = 1.16e-4. Overall: Spearman’s rho = -0.62 p = 2.3e-4.

### Several haplotypes were exclusive to the mother or the infant

Parasite genetics may play a role in the probability of congenital transmission of *T. cruzi*. To address this possibility, we searched for haplotypes that were more or less likely to be transmitted from the mother to her infant. Haplotypes found in a mother, but not her infant, could represent clones that are less likely to be congenitally transmitted. For this analysis, we counted the number of times a haplotype was found in a mother, her paired infant, or in both the mother and her infant (Fig 3a). Haplotypes 0, 1 and 7 appeared in both samples of some mother-infant pairs, indicating that these haplotypes were likely to be transmitted and detected in maternal blood. Haplotypes 3, 5, and 6 were only found in mothers and not paired infants. Haplotypes 4 and 9 were only ever found in the infant samples of mother-infant pairs. To further explore the effect of haplotype on transmission, we also analyzed haplotype presence in samples without considering the mother-infant pairs (Fig 3b). Among these samples, haplotypes 5 and 6 were no longer exclusive to the mother and were found in some infant samples, while haplotype 3 remained exclusive to mothers. Interestingly, haplotypes 4 and 9 remain exclusive to infant samples. This indicates that these haplotypes are highly likely to be transmitted, and that there may be additional biological mechanisms resulting in the lack of these haplotype’s detection in the maternal blood at time of birth.

**Figure 3:**
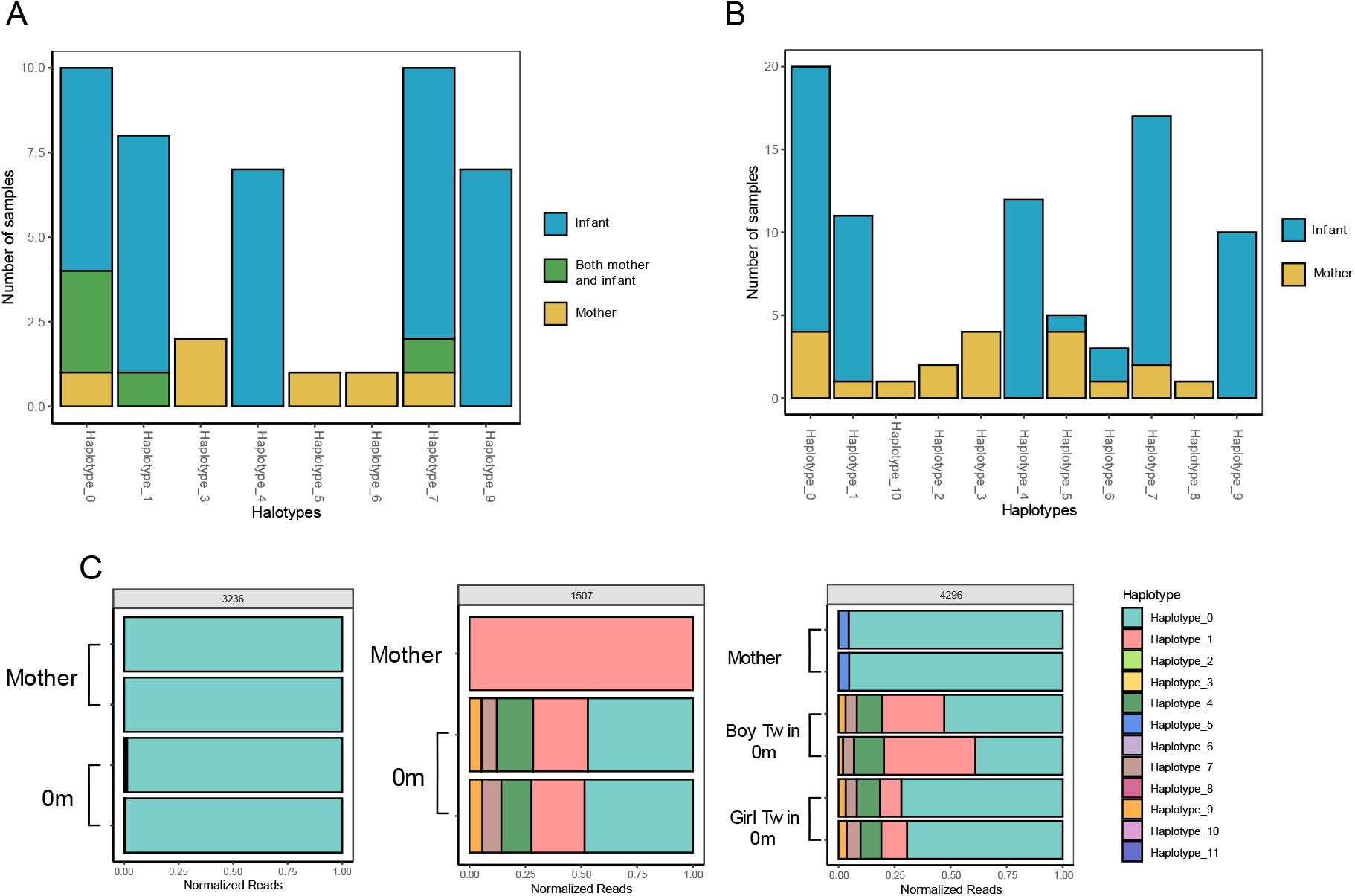
Certain haplotypes are found only in mothers or only in infants. **A**. Among families with paired infant-mother samples, the number of times a haplotype was found only in the mother sample, only in the newborn sample, or in both the mother and the newborn sample. **B**. Number of times a haplotype was found in any mother or any newborn, regardless of family, including samples without mother-infant pairs. **C**. Representative plots of relative haplotype abundance in each sample grouped by family. Read count was normalized to total reads in each sample.

Examining individual families reveals diverse transmission patterns (Fig 3c, Supplemental figure 2, and 3). Family 1507 shows the maternal detection of haplotype 1, while within the infant sample we detect haplotype 1 in addition to several other haplotypes, consistent with our previous observation of increased diversity in infant samples relative to mother’s samples. Transmission within family 4296 is perhaps the most informative. In this case, two haplotypes are detected in the maternal sample, while a larger number of haplotypes are detected in each of the two fraternal twins. Notably, the haplotype distribution in each twin is nearly identical. Because these were fraternal twins, with two separate placentas, this finding supports the hypothesis that transmission is not the result of a single inoculating incident.

## Discussion

These data describe the changes in parasite diversity that exist during the process of *T. cruzi* congenital transmission and suggest a role for parasite genetics in the likelihood of transmission. By directly sequencing blood samples from patients and targeting a single copy gene by amplicon sequencing, we were able to identify complex infections in mother and infant samples in an endemic setting and determine changes in parasite strain diversity that occur during the process of transmission. While our findings are somewhat limited by small sample sizes, our approach allows a direct assessment of parasite diversity beyond simple DTU classification and reveals new features of congenital transmission of *T. cruzi* that warrant further investigation.

The unique method used in this paper allows us to avoid over-estimating complexity of infection for a sample. Here we assume the single locus gene occurs twice per parasite, with a unique allele on each homologous chromosome, thereby potentially underestimating the parasite diversity. It must be stated that this assumption of exclusive diploidy may not always hold; karyotypic instability has been found in *T. cruzi*, and gene duplication occurs commonly (19,20). However, duplicated genes often encode surface genes involved in immune evasion, and the conserved metabolic gene targeted in this work is not thought to be expressed on the surface. An additional strength of this work is that parasites were sequenced directly from patient samples and did not undergo expansion and potential strain selection in culture. Thus, we avoid the possibility of selecting for clones that are better adapted to culture at the expense of the true parasite diversity. This study demonstrates the feasibility of this approach for characterizing parasite diversity across congenital infection.

The majority of the haplotypes detected in our samples were hybrid DTUs, likely TcV considering other strain typing work from the same setting (21). In this study, we observe potential ancestral recombination events between the parental TcII and TcIII alleles in our recovered haplotypes. This suggests additional diversity exists in the hybrid strains beyond what has been sequenced in the reference CL Brener strain. Within these hybrid DTU types, we observed several haplotypes that occurred exclusively in mother or infant samples. This suggests that there may be genetic factors within DTUs that influence parasite transmission and underscores the fact that current DTU designations are insufficient to appropriately assess the diversity of parasite strains in a single infection. Investigation into specific virulence factors, either by transcriptomic or genomic analysis, rather than DTU typing, is likely to be required to uncover the factors that influence transmission. An analysis including mothers infected with a more diverse set of parasite DTUs and mothers that did not transmit to their infant may shed light on the degree to which DTU alone can influence the probability of transmission.

While parasite genetics may influence the probability of transmission, our data suggest that there is no significant bottleneck at transmission, with complex infections observed in both mother and infant samples. Surprisingly, we observe an increase in parasite diversity after transmission, with many haplotypes recovered in the infant that were undetectable in maternal blood. This finding is corroborated in a similar study, where Lewellyn *et al* found novel strain types in infants compared to their paired mothers using a diverse low copy number gene (22). In that study, it was unclear whether the observed “strain types” represented individual parasite clones or diversity that was generated during infection, as the locus analyzed was a variant surface protein that is likely to diversify during the course of infection. Because our approach analyzes a single-copy metabolic gene and most haplotypes were identified independently in multiple infections, we are able to more confidently assume that detected haplotypes were not the result of diversification during infection.

The increased diversity in the infant samples may be explained in several ways. Perhaps most intriguing is the possibility that some parasite clones prefer the placental environment, remaining undetectable in the maternal blood. If these *T. cruzi* clones are sequestered in the placenta, they might not circulate at detectable levels in the mother’s blood. A previous study performed kDNA PCR on Chagas positive mothers and found multiple patients with negative bloodstream but positive placental PCR, and in two cases found placental minicircle fragments that were not detectable in the bloodstream of the same patient, suggesting placental tropism for specific parasite clones (23). Placental tropism of specific *T. cruzi* strains has also been observed in mouse models of infection (24,25).

An additional explanation is that parasite transmission occurs throughout pregnancy. If parasite clones cross the placenta during multiple waves of infection, the composition in the circulation of the infant would reflect the set of parasite clones present in the mother throughout pregnancy, even if these clones are absent from the mother’s circulation at the time of delivery. It is possible that waves of parasitemia lead to the expansion of different parasite clones at different time points in the maternal blood during pregnancy, and that clones found in the maternal blood at birth may not represent the true diversity of parasite clones infecting the mother.

While compelling biological explanations for this observed increase in diversity exist, it remains possible that this observation is simply a matter of sampling. The lower parasitemia of chronically infected mothers means that fewer parasites are sampled from mothers for sequencing, potentially biasing diversity estimates. While our other analyses suggest that mother samples are not undersampled, a correlation between parasitemia and Shannon’s diversity means we cannot preclude this possibility.

Importantly, however, the observation of complex infections in infants is not affected by these issues. If infants are infected with multiple clones, regardless of each clone’s presence in the mother, this is most likely the result of several parasites successfully colonizing the infant. Little is known about the dynamics of congenital transmission of *T. cruzi*, but the relatively low rate of congenital transmission compared to other parasitic congenital infection such as toxoplasmosis (∼5% for Chagas compared to as high as 65% for acute toxoplasmosis (26)) might suggest that it occurs as a result of a single parasite occasionally breaching the placental barrier. Contrary to this model, our data suggest that congenital infection is the result of several parasites infecting the infant. The parasite profile from fraternal twin samples, distinct from the mother but identical between siblings, supports this model of transmission. Each of the two placentas were infected with the same set of clones, probably during multiple independent infection events during their gestation. The temporal dynamics of this process remain unclear. Parasites may cross the placenta at certain times during infection, which is in line with previous reports showing that parasitemia during the third trimester is most predictive of congenital infection (23). Alternatively, parasite clones might cross the placenta during multiple waves of infection.

Why transmission of multiple parasites occurs in some pregnancies, while most others result in no infection, will be a central question going forward. The fact that multiple parasites cross the placenta suggests that host factors, including the integrity of the placental barrier and the potency of the maternal immune response, likely influence the probability of transmission.

This study defines the changes in parasite diversity that occur during congenital transmission and raises interesting questions about the mechanism of the process. We find no evidence of a transmission bottleneck during congenital infection, which, together with haplotype data from the fraternal twins in this study, support a model whereby multiple parasites colonize the infant during pregnancy. Moreover, we detect two haplotypes unique to newborn samples that were not detected in maternal peripheral blood, which together with previously published data suggest a link between parasite genetics and transmission probability (6). Understanding the mechanisms influencing transmission of *T. cruzi* may help inform better diagnostics and lead to more effective treatment, limiting the global burden of Chagas disease.

