## Supplemental methods for "Amplicon sequencing reveals complex infection in infants congenitally infected with *Trypanosoma cruzi* and informs the dynamics of parasite transmission"

Supplementary methods

T. cruzi nested PCR amplicons were prepared for sequencing using first round primers Tcs5dFOR 5’- GGACGTGGCGTTTGATTTAT-3’ and Tcsc5dREV 5’- TCCCATCTTCTTCGTTGACT -3’ and cycling conditions: 98°C x 30 sec; 40 cycles of 98°C x 10s, 55°C x 30s, 72°C x 30 sec; and 72°C x 10 min. Amplification was performed on a Bio-Rad T100 Thermal Cycler (Bio-Rad, Hercules, California) in 20 µL reactions containing 5 µL 10x Phusion Buffer (Roche, Madison, WI), 0.3 ul Phusion Taq, 0.5 ul (20 uM) Forward and Reverse primers, 0.4 µL 10 mM dNTPs, and 2 µL template DNA. First round PCR products were diluted 1:10 with water and then 2 mL was used as template DNA in the second round PCR using primers, modified by inclusion of a multiplex identifier (MID) sequence, listed in Supplementary Methods. Second round of nested PCR assay was set up as follows: 2 µL of template DNA (diluted 1:10) PCR product from R1, 2.5 µL of 10X MgCl2 buffer, 0.5 µL (20 uM) forward and reverse primer, each tagged with Roche multiplex identifiers (MIDs, listed below), 0.5 µL (10 mM) dNTPs (New England BioLabs, catalog number N0447L), 0.25 µL Roche tag and 18.75 µL nuclease-free water. The PCR conditions were as follows: initial denaturation (94°C for 2 minutes), followed by 40 cycles of denaturation (94°C for 60 seconds), annealing (55°C for 30 seconds), extension (72°C for 60 seconds), and final extension (72°C for 5 minutes). PCR products were visualized on 1% (w/v) agarose (Sigma-Aldrich, catalog number A9539-500G) gels stained with RedSafeNucleic Acid Staining Solution (iNtRON Biotechnology DR, catalog number 21141). If amplification failed, the cycle number in both rounds was increased to 45 cycles.

Library Preparation and Sequencing PCR amplicons were purified using the Qiagen PCR purification Kit (Qiagen catalog number 28104) according to the manufacturer’s instructions. The purified products were quantified using Quant-iT dsDNA Assay Kit, High Sensitivity (Thermo Fisher, catalog number Q33120) according to the manufacturer’s instructions. The quantified PCR amplicons were normalized by diluting in EB buffer to achieve equimolar concentrations of 50ng each across all samples and thereafter mixed to create amplicon pools of nonoverlapping MIDs. Sequence library preparation of the amplicon pools was performed according to the manufacturer’s instructions using the KAPA HyperPrep Kit (Roche, catalog number KK8504) with BioScientific index adapters. Successful library preparation was confirmed using a random sample of 10 libraries on the Agilent High Sensitivity D1000 ScreenTape System (Agilent, catalog number 5067-5584). Adapter-ligated amplicon libraries were then quantified, normalized to 1 ng each, and mixed to create final pool. Paired-end sequencing (2 × 300 bp chemistry) of the final pool was performed on the Illumina MiSeq platform using the MiSeq Reagent Kit v3 (Illumina, catalog MS-102-3003 number) at the University of North Carolina at Chapel Hill high-throughput sequencing facility.

Barcoded forward and reverse primers for 2^nd^ PCR

TcSc5D_308F_MID1 ACGAGTGCGTTCACCGTGGGTACAGTAAAAT

TcSc5D_308F_MID2 ACGCTCGACATCACCGTGGGTACAGTAAAAT

TcSc5D_308F_MID3 AGACGCACTCTCACCGTGGGTACAGTAAAAT

TcSc5D_308F_MID4 AGCACTGTAGTCACCGTGGGTACAGTAAAAT

TcSc5D_308F_MID6 ATATCGCGAGTCACCGTGGGTACAGTAAAAT

TcSc5D_308F_MID7 CGTGTCTCTATCACCGTGGGTACAGTAAAAT

TcSc5D_308F_MID8 CTCGCGTGTCTCACCGTGGGTACAGTAAAAT

TcSc5D_308F_MID10 TCTCTATGCGTCACCGTGGGTACAGTAAAAT

TcSc5D_308F_MID11 TGATACGTCTTCACCGTGGGTACAGTAAAAT

TcSc5D_308F_MID13 CATAGTAGTGTCACCGTGGGTACAGTAAAAT

TcSc5D_308F_MID14 CGAGAGATACTCACCGTGGGTACAGTAAAAT

TcSc5D_308F_MID15 ATACGACGTATCACCGTGGGTACAGTAAAAT

TcSc5D_308F_MID16 TCACGTACTATCACCGTGGGTACAGTAAAAT

TcSc5D_308F_MID17 CGTCTAGTACTCACCGTGGGTACAGTAAAAT

TcSc5D_308F_MID18 TCTACGTAGCTCACCGTGGGTACAGTAAAAT

TcSc5D_308F_MID_19 TGTACTACTCTCACCGTGGGTACAGTAAAAT

TcSc5D_308F_MID_20 ACGACTACAGTCACCGTGGGTACAGTAAAAT

TcSc5D_308F_MID_21 CGTAGACTAGTCACCGTGGGTACAGTAAAAT

TcSc5D_308F_MID-22 TACGAGTATGTCACCGTGGGTACAGTAAAAT

TcSc5D_308F_MID_23 TACTCTCGTGTCACCGTGGGTACAGTAAAAT

TcSc5D_308F_MID_24 TAGAGACGAGTCACCGTGGGTACAGTAAAAT

TcSc5D_308F_MID_25 TCGTCGCTCGTCACCGTGGGTACAGTAAAAT

TCSC5D_Rev_MID1 ACGCACTCGTTCCCATCTTCGTTGACT

TCSC5D_Rev_MID2 TGTCGAGCGTTCCCATCTTCGTTGACT

TCSC5D_Rev_MID3 GAGTGCGTCTTCCCATCTTCGTTGACT

TCSC5D_Rev_MID4 CTACAGTGCTTCCCATCTTCGTTGACT

TCSC5D_Rev_MID6 CTCGCGATATTCCCATCTTCGTTGACT

TCSC5D_Rev_MID7 TAGAGACACGTCCCATCTTCGTTGACT

TCSC5D_Rev_MID8 GACACGCGAGTCCCATCTTCGTTGACT

TCSC5D_Rev_MID10 CGCATAGAGATCCCATCTTCGTTGACT

TCSC5D_Rev_MID11 AGACGTATCATCCCATCTTCGTTGACT

TCSC5D_Rev_MID13 CACTACTATGTCCCATCTTCGTTGACT

TCSC5D_Rev_MID14 GTATCTCTCGTCCCATCTTCGTTGACT

TCSC5D_Rev_MID15 TACGTCGTATTCCCATCTTCGTTGACT

TCSC5D_Rev_MID16 TAGTACGTGATCCCATCTTCGTTGACT

TCSC5D_Rev_MID17 GTACTAGACGTCCCATCTTCGTTGACT

TCSC5D_Rev_MID18 GCTACGTAGATCCCATCTTCGTTGACT

TCSC5D_Rev_MID19 GAGTAGTACATCCCATCTTCGTTGACT

TCSC5D_Rev_MID20 CTGTAGTCGTTCCCATCTTCGTTGACT

TCSC5D_Rev_MID21 CTAGTCTACGTCCCATCTTCGTTGACT

TCSC5D_Rev_MID22 CATACTCGTATCCCATCTTCGTTGACT

TCSC5D_Rev_MID23 CACGAGAGTATCCCATCTTCGTTGACT

TCSC5D_Rev_MID24 CTCGTCTCTATCCCATCTTCGTTGACT

TCSC5D_Rev_MID25 CGAGCGACGATCCCATCTTCGTTGACT
