## Supplemental data for "Amplicon sequencing reveals complex infection in infants congenitally infected with *Trypanosoma cruzi* and informs the dynamics of parasite transmission"

>Haplotype_0

ATATTACAATGTTTCTGATTATGGATGGCCGTATCTTTTTCTTAGTATATTGATGTTTTTTATCTTTACGGACTTTATGGTGTATTGGTTTCATCGTGGTTTACATCACCCAACATTATACCGATACCTTCATAAATTACATCATACATACAAATATACAACACCATTTTCATCTCATGCATTTAATCCTTGTGATGGATTTGGTCAAGGTTCACCATATTACGCATTTATTTTTTTATTTCCTATGCATAATTACCTTTTTGTTATTCTCTTTTTCGCTGTTAACCTATGGACAATTTCTATTCATGATCAAGTAGACTTTGGAGGTCATTTTGTGAACACAACAGGACACCATACAATTCATCATGTACTTTTTAATTACGACTATGGACAGTACTTTACCGTATGGGACCGTATTGGCGGAACTTATAAACCCGCACAACAGACGCATCTTTTTCCATTATTTACAAAAGGTGGACGCATTGAAGAGGTTG

>Haplotype_1

ATATTACAATGTTTCCGATTATGGATGGTCATATCTTTTTTTTAGTATATTGATGTTTTTTATTTTTACGGATTTTATGGTGTATTGGTTTCATCGTGGTTTACATCATCCAACATTATACCGATATCTTCATAAATTACATCATACATACAAATATACCACACCATTTTCATCGCATGCATTTAATCCTTGTGATGGATTTGGTCAAGGTTCACCATATTACGCATTTATTTTTTTATTTCCGATGCATAATTATCTTTTTGTGATTCTCTTTTTTGCCGTTAATTTATGGACCATCTCTATTCATGATCAGGTGGATTTTGGAGGGCATTTTGTCAACACAACCGGACATCATACAATTCATCATGTACTTTTTAATTACGACTATGGACAGTACTTTACCGTATGGGATCGTATTGGTGGAACATATAAACCGGCACAACAGACACATCATTTCCCATTATTTACAAAAGGTGGACGCAATGAAGAGGTTG

>Haplotype_2

ATATTACAATGTTTCTGATTATGGATGGCCGTATCTTTTTCTTAGTATATTGATGTTTTTTATCTTTACGGACTTTATGGTATATTGGTTTCATCGTGGTTTACATCACCCAACATTATACCGATACCTTCATAAATTACATCATACATACAAATATACAACACCATTTTCATCTCATGCATTTAATCCTTGTGATGGATTTGGTCAAGGTTCACCATATTACGCATTTATTTTTTTATTTCCTATGCATAATTACCTTTTTGTTATTCTCTTTTTCGCTGTTAACCTATGGACAATCTCTATTCATGATCAAGTAGACTTTGGAGGTCATTTTGTGAACACAACAGGACACCATACAATTCATCATGTACTTTTTAATTACGACTATGGACAGTACTTTACCGTATGGGATCGTATTGGTGGAACATATAAACCGGCACAACAGACACATCATTTCCCATTATTTACAAAAGGTGGACGCAATGAAGAGGTTG

>Haplotype_3

ATATTACAATGTTTCTGATTATGGATGGCCGTATCTTTTTCTTAGTATATTGATGTTTTTTATCTTTACGGACTTTATGGTGTATTGGTTTCATCGTGGTTTACATCACCCAACATTATACCGATACCTTCATAAATTACATCATACATACAAATATACAACACCATTTTCATCTCATGCATTTAATCCTTGTGATGGATTTGGTCAAGGTTCACCATATTACGCATTTATTTTTTTATTTCCTATGCATAATTACCTTTTTGTTATTCTCTTTTTCGCTGTTAACCTATGGACAATTTCTATTCATGATCAAGTAGACTTTGGAGGTCATTTTGTGAACACAACAGGACACCATACAATTCATCATGTACTTTTTAATTACGACTATGGACAGTACTTTACCGTATTGGACCGTATTGGCGGAACTTATAAACCCGCACAACAGACGCATCTTTTTCCATTATTTACAAAAGGTGGACGCATTGAAGAGGTTG

>Haplotype_4

ATATTACAATGTTTCTGATTATGGATGGCCGTATCTTTTTCTTAGTATATTGATGTTTTTTATCTTTACGGACTTTATGGTGTATTGGTTTCATCGTGGTTTACATCACCCAACATTATACCGATACCTTCATAAATTACATCATACATACAAATATACAACACCATTTTCATCTCATGCATTTAATCCTTGTGATGGATTTGGTCAAGGTTCACCATATTACGCATTTATTTTTTTATTTCCGATGCATAATTATCTTTTTGTGATTCTCTTTTTTGCCGTTAATTTATGGACCATCTCTATTCATGATCAGGTGGATTTTGGAGGGCATTTTGTCAACACAACCGGACATCATACAATTCATCATGTACTTTTTAATTACGACTATGGACAGTACTTTACCGTATGGGATCGTATTGGTGGAACATATAAACCGGCACAACAGACACATCATTTCCCATTATTTACAAAAGGTGGACGCAATGAAGAGGTTG

>Haplotype_5

ATATTACAATGTTTCTGATTATGGATGGCCGTATCTTTTTCTTAGTATATTGATGTTTTTTATCTTTACGGACTTTATGGTATATTGGTTTCATCGTGGTTTACATCACCCAACATTATACCGATACCTTCATAAATTACATCATACATACAAATATACAACACCATTTTCATCTCATGCATTTAATCCTTGTGATGGATTTGGTCAAGGTTCACCATATTACGCATTTATTTTTTTATTTCCTATGCATAATTACCTTTTTGTTATTCTCTTTTTCGCTGTTAACCTATGGACAATCTCTATTCATGATCAAGTAGACTTTGGAGGTCATTTTGTGAACACAACAGGACACCATACAATTCATCATGTACTTTTTAATTACGACTATGGACAGTACTTTACCGTATGGGACCGTATTGGCGGAACTTATAAACCCGCACAACAGACGCATCTTTTCCCATTATTTACAAAAGGTGGACGCATTGAAGAGGTTG

>Haplotype_12

ATATTACAATGTTTCTGATTATGGATGGCCGTATCTTTTTCTTAGTATATTGATGTTTTTTATCTTTACGGACTTTATGGTGTATTGGTTTCATCGTGGTTTACATCACCCAACATTATACCGATACCTTCATAAATTACATCATACATACAAATATACAACACCATTTTCATCTCATGCATTTAATCCTTGTGATGGATTTGGTCAAGGTTCACCATATTACGCATTTATTTTTTTATTTCCTATGCATAATTACCTTTTTGTTATTCTCTTTTTCGCTGTTAACCTATGGACAATTTCTATTCATGATCAAGTAGACTTTGGAGGTCATTTTGTGAACACAACAGGACACCATACAATTCATCATGTACTTTTTAATTACGACTATGGACAGTACTTTACCGTATGGGACCGTATTGGCGGAACTTATAAACCCGCACAACAGACGCATCTTTTTCCATTATTTACAAAAGGTGGACGCATTGAAGAGGTTGAGTCAACGAAGAAGATGGG

>Haplotype_7

ATATTACAATGTTTCTGATTATGGATGGCCGTATCTTTTTCTTAGTATATTGATGTTTTTTATCTTTACGGACTTTATGGTGTATTGGTTTCATCGTGGTTTACATCACCCAACATTATACCGATACCTTCATAAATTACATCATACATACAAATATACAACACCATTTTCATCTCATGCATTTAATCCTTGTGATGGATTTGGTCAAGGTTCACCATATTACGCATTTATTTTTTTATTTCCTATGCATAATTACCTTTTTGTTATTCTCTTTTTCGCTGTTAACCTATGGACAATTTCTATTCATGATCAAGTAGACTTTGGAGGTCATTTTGTGAACACAACAGGACACCATACAATTCATCATGTACTTTTTAATTACGACTATGGACAGTACTTTACCGTATGGGATCGTATTGGTGGAACATATAAACCGGCACAACAGACACATCATTTCCCATTATTTACAAAAGGTGGACGCAATGAAGAGGTTG

>Haplotype_8

ATATTACAATGTTTCTGATTATGGATGGCCGTATCTTTTTCTTAGTATATTGATGTTTTTTATCTTTACGGACTTTATGGTGTATTGGTTTCATCGTGGTTTACATCACCCAACATTATACCGATACCTTCATAAATTACATCATACATACAAATATACAACACCATTTTCATCTCATGCATTTAATCCTTGTGATGGATTTGGTCAAGGTTCACCATATTACGCATTTATTTTTTTATTTCCTATGCATAATTACCTTTTTGCTATTCTCTTTTTCGCTGTTAACCTATGGACAATTTCTATTCATGATCAAGTAGACTTTGGAGGTCATTTTGTGAACACAACAGGACACCATACAATTCATCATGTACTTTTTAATTACGACTATGGACAGTACTTTACCGTATGGGACCGTATTGGCGGAACTTATAAACCCGCACAACAGACGCATCTTTTTCCATTATTTACAAAAGGTGGACGCATTGAAGAGGTTG

>Haplotype_9

ATATTACAATGTTTCTGATTATGGATGGCCGTATCTTTTTCTTAGTATATTGATGTTTTTTATCTTTACGGACTTTATGGTGTATTGGTTTCATCGTGGTTTACATCACCCAACATTATACCGATACCTTCATAAATTACATCATACATACAAATATACAACACCATTTTCATCTCATGCATTTAATCCTTGTGATGGATTTGGTCAAGGTTCACCATATTACGCATTTATTTTTTTATTTCCGATGCATAATTATCTTTTTGTGATTCTCTTTTTTGCCGTTAATTTATGGACCATCTCTATTCATGATCAGGTGGATTTTGGAGGGCATTTTGTCAACACAACCGGACATCATACAATTCATCATGTACTTTTTAATTACGACTATGGACAGTACTTTACCGTATGGGACCGTATTGGCGGAACTTATAAACCCGCACAACAGACGCATCTTTTTCCATTATTTACAAAAGGTGGACGCATTGAAGAGGTTG

>Haplotype_10

ATATTACAATGTTTCTGATTATGGATGGCCGTATCTTTTTCTTAGTATATTGATGTTTTTTATCTTTACGGACTTTATGGTATATTGGTTTCATCGTGGTTTACATCATCCAACATTATACCGATACCTTCATAAATTACATCATACATACAAATATACAACACCATTTTCATCTCATGCATTTAATCCTTGTGATGGATTTGGTCAAGGTTCACCATATTACGCATTTATTTTTTTATTTCCGATGCATAATTACCTTTTTGTTATTCTCTTTTTCGCTGTTAACCTATGGACAATCTCTATTCATGATCAAGTAGACTTTGGAGGTCATTTTGTGAACACAACAGGACACCATACAATTCATCATGTACTTTTTAATTACGACTATGGACAGTACTTTACCGTATGGGACCGTATTGGCGGAACTTATAAACCCGCACAACAGACGCATCTTTTCCCATTATTTACAAAAGGTGGACGCAATGAAGAGGTTG

>Haplotype_11

ATATTACAATGTTTCTGATTATGGATGGCCGTATCTTTTTCTTAGTATATTGATGTTTTTTATCTTTACGGACTTTATGGTATATTGGTTTCATCGTGGTTTACATCACCCAACATTATACCGATACCTTCATAAATTACATCATACATACAAATATACAACACCATTTTCATCTCATGCATTTAATCCTTGTGATGGATTTGGTCAAGGTTCACCATATTACGCATTTATTTTTTTATTTCCTATGCATAATTACCTTTTTGTTATTCTCTTTTTCGCTGTTAACCTATGGACAATCTCTATTCATGATCAAGTAGACTTTGGAGGTCATTTTGTGAACACAACAGGACACCATACAATTCATCATGTACTTTTTAATTACGACTATGGACAGTACTTTACCGTATGGGACCATATTGGCGGAACTTATAAACCCGCACAACAGACGCATCTTTTCCCATTATTTACAAAAGGTGGACGCATTGAAGAGGTTG

>Haplotype_6

ATATTACAATGTTTCCGATTATGGATGGTCATATCTTTTTTTAAGTATTTTGATGTTTTTTATCTTTACGGATTTTATGGTTTATTGGTTTCATCGTGGTTTACATCATCCCACATTATACCGATATCTTCATAAATTACATCATACATACAAATATACCACACCATTTTCATCTCATGCATTTAATCCTTGTGATGGATTTGGTCAAGGTTCACCATATTATGCATTTATTTTTTTATTTCCTATGCATAATTATCTTTTTGTTATTCTCTTTTTTGCCGTCAATTTATGGACCATCTCCATTCACGATCAGGTGGATTTTGGAGGGCATTTTGTTAACACAACCGGACATCATACAATTCATCATGTACTTTTTAATTACGACTACGGACAATACTTCACCGTATGGGATCGTATTGGTGGAACGTATAAACCGGCACAACAGACGCATCATTTCCCGTTATTTACAAAAGGTGGTCGTAATGAAGAGATGA
