## Supplementary figures and images for "Amplicon sequencing reveals complex infection in infants congenitally infected with *Trypanosoma cruzi* and informs the dynamics of parasite transmission"

### Supplemental Figure 1

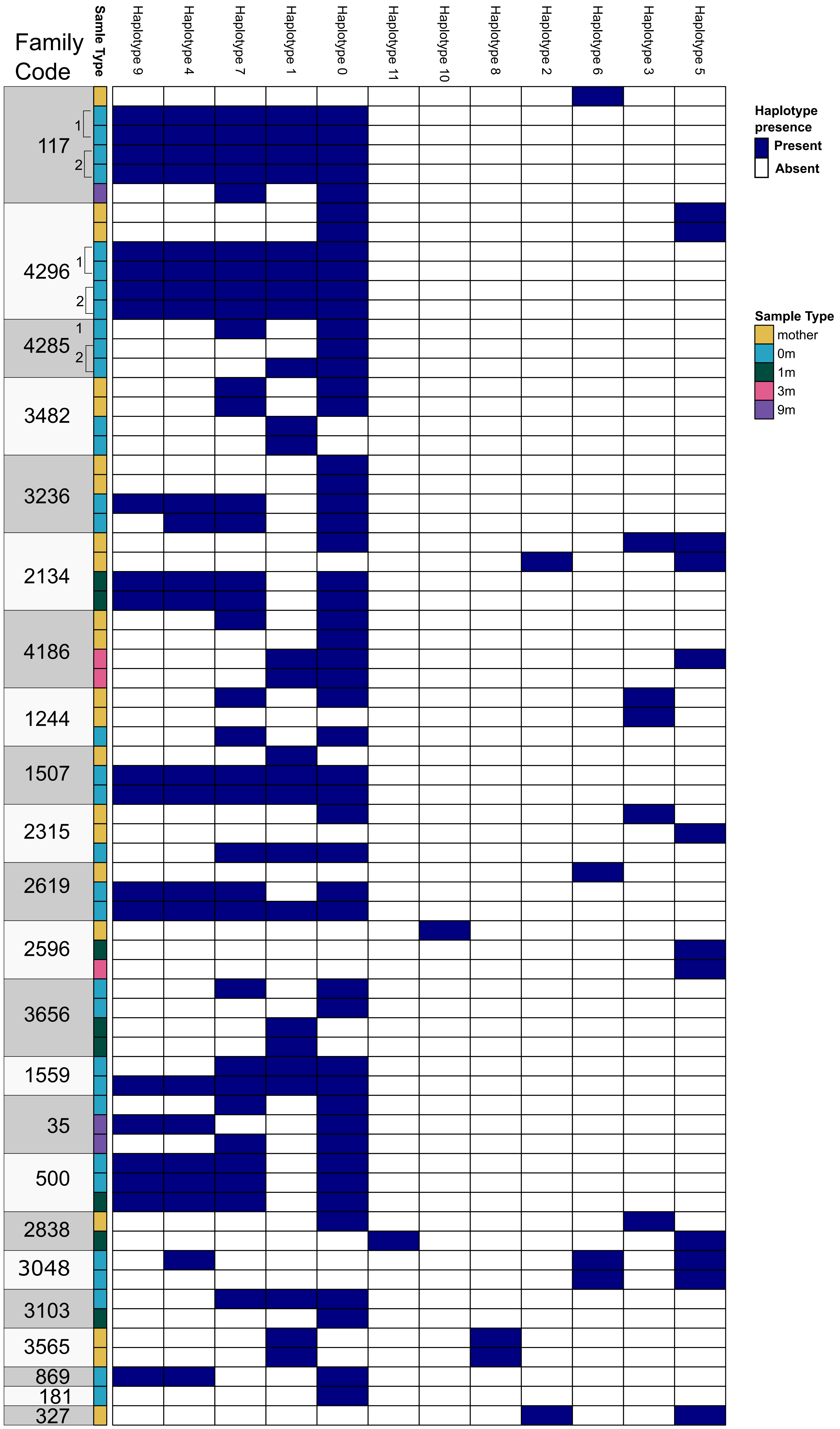

### Supplemental Figure 2

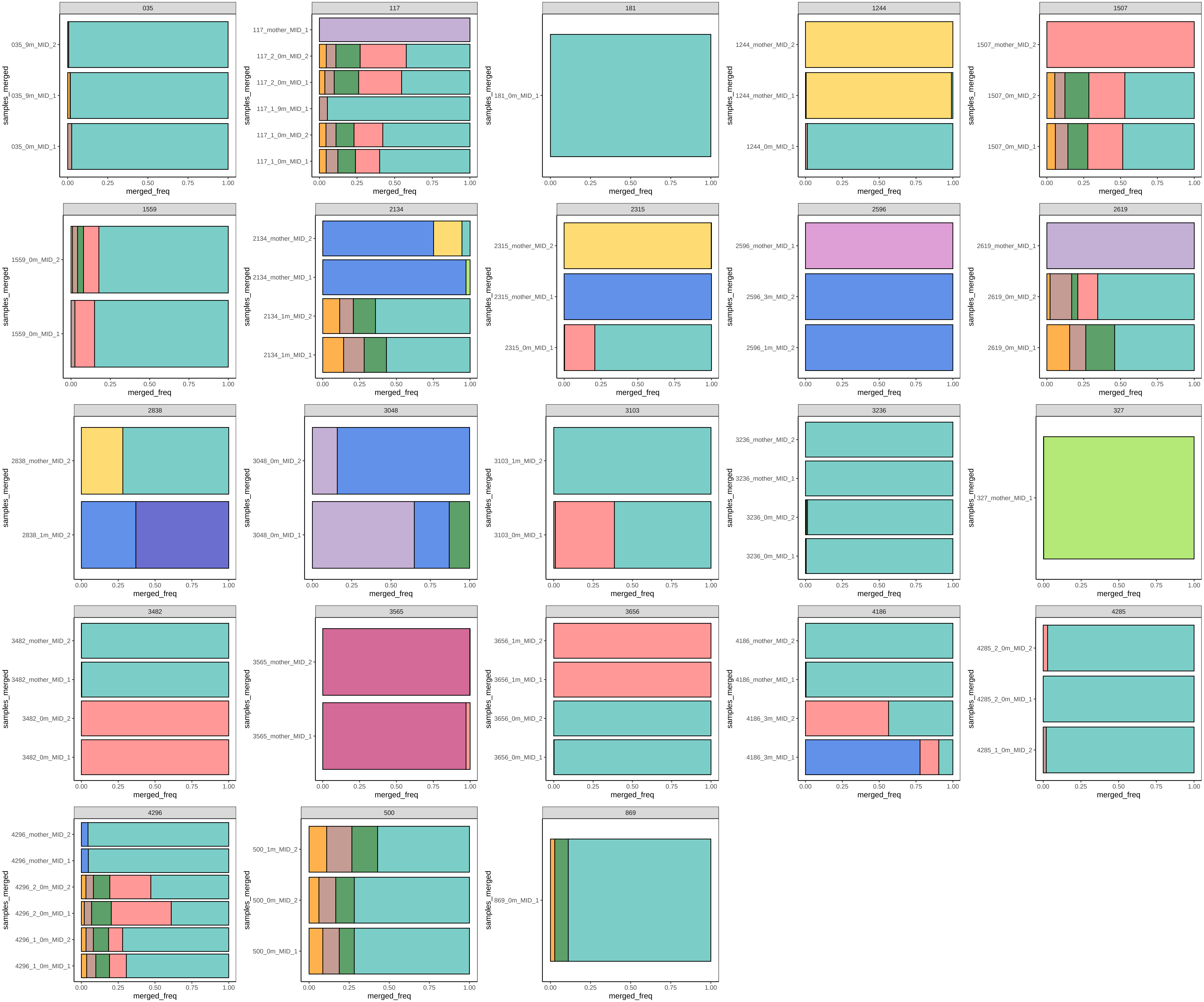

### Supplemental Figure 3

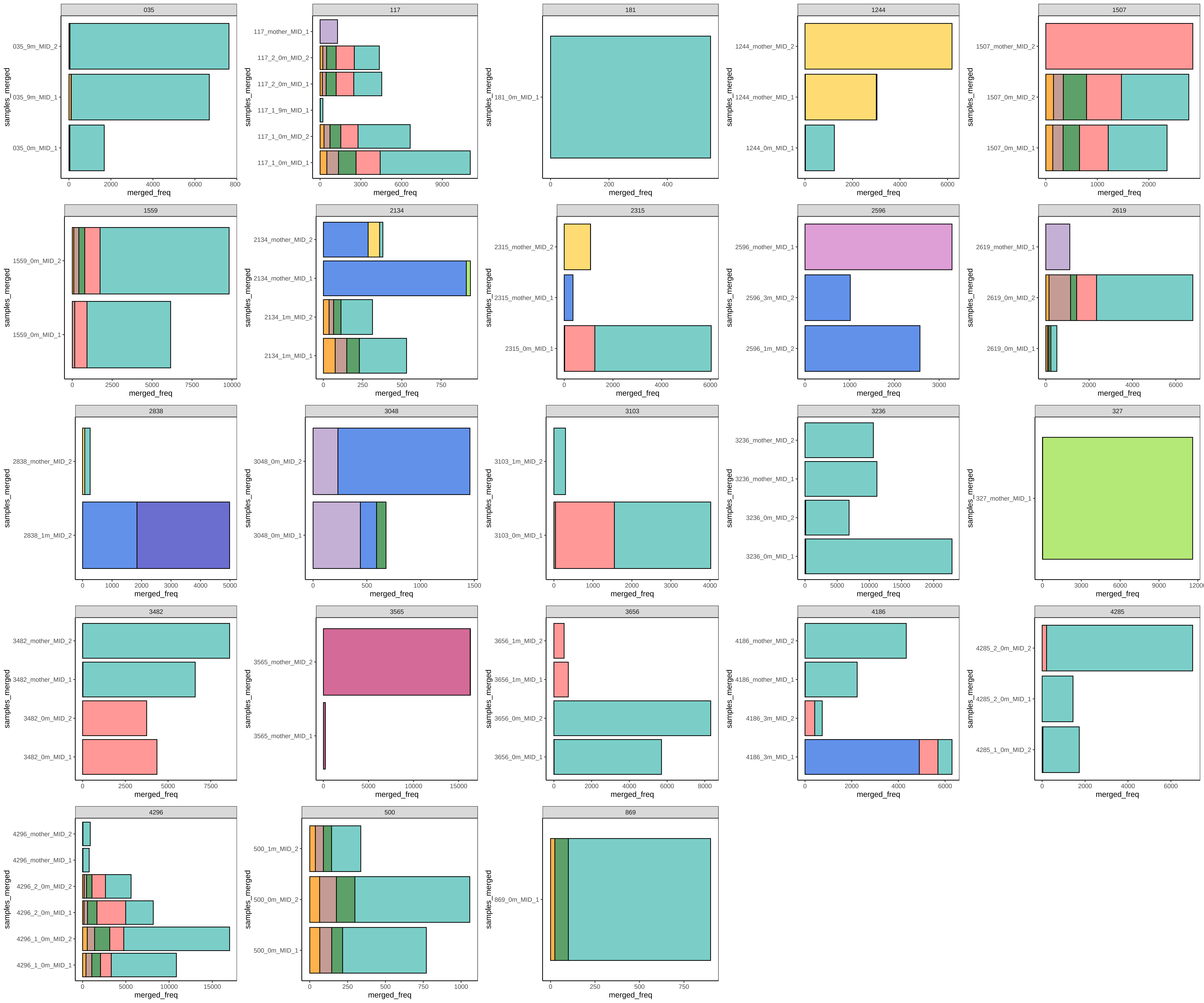
