## Supplementary material for "Amplicon sequencing reveals complex infection in infants congenitally infected with *Trypanosoma cruzi* and informs the dynamics of parasite transmission": Figure Legends

**Supplemental Figure 1: Up to six haplotypes could be detected in a single sample**. Heatmap showing the haplotypes detected in each sample. Each column is a haplotype while rows are each sample. Samples are grouped by family and annotation bar indicates sample type. All samples were run in duplicate, though samples with insufficient read depth were excluded. Families 117, 4296 and 4285 included twins, which are enumerated each 1 and 2.

**Supplemental Figure 2: Haplotype abundance in each family normalized to read depth.** All plots of relative haplotype abundance in each sample grouped by family. Read count was normalized to total reads in each sample. Number following MID indicates replicate.

**Supplemental Figure 3: Haplotype abundance in each family measured by total number of reads.** All plots of relative haplotype abundance in each sample grouped by family. Read count was not normalized and each x axis represents the total number of reads each called haplotype received. MID number indicates replicate. Number following MID indicates replicate.
